# Population Replacement in Early Neolithic Britain

**DOI:** 10.1101/267443

**Authors:** Selina Brace, Yoan Diekmann, Thomas J. Booth, Zuzana Faltyskova, Nadin Rohland, Swapan Mallick, Matthew Ferry, Megan Michel, Jonas Oppenheimer, Nasreen Broomandkhoshbacht, Kristin Stewardson, Susan Walsh, Manfred Kayser, Rick Schulting, Oliver E. Craig, Alison Sheridan, Mike Parker Pearson, Chris Stringer, David Reich, Mark G. Thomas, Ian Barnes

**Affiliations:** Department of Earth Sciences, Natural History Museum, London SW7 5BD, UK; Research Department of Genetics, Evolution and Environment, University College London, London WC1E 6BT, UK; Department of Genetics, Harvard Medical School, Boston, Massachusetts 02115, USA; Broad Institute of MIT and Harvard, Cambridge, Massachusetts 02142, USA; Howard Hughes Medical Institute, Harvard Medical School, Boston, Massachusetts 02115, USA; Department of Biology, Indiana University Purdue University, Indianapolis (IUPUI), Indianapolis, IN, USA; Department of Genetic Identification, Erasmus MC University Medical Centre Rotterdam, Rotterdam, The Netherlands; Institute of Archaeology, University of Oxford, 36 Beaumont St, Oxford,OX1 2PG; BioArCh, Department of Archaeology, University of York, UK; National Museums Scotland, Edinburgh EH1 1JF, UK; Institute of Archaeology, University College London,London WC1H 0PY, UK

**Author notes:** these authors contributed equally (Selina Brace, Yoan Diekmann, Thomas J. Booth). these authors co-supervised the work (David Reich, Mark G. Thomas, Ian Barnes).

## Abstract

The roles of migration, admixture and acculturation in the European transition to farming have been debated for over 100 years. Genome-wide ancient DNA studies indicate predominantly Anatolian ancestry for continental Neolithic farmers, but also variable admixture with local Mesolithic hunter-gatherers^1–9^. Neolithic cultures first appear in Britain c. 6000 years ago (kBP), a millennium after they appear in adjacent areas of northwestern continental Europe. However, the pattern and process of the British Neolithic transition remains unclear^10–15^. We assembled genome-wide data from six Mesolithic and 67 Neolithic individuals found in Britain, dating from 10.5-4.5 kBP, a dataset that includes 22 newly reported individuals and the first genomic data from British Mesolithic hunter-gatherers. Our analyses reveals persistent genetic affinities between Mesolithic British and Western European hunter-gatherers over a period spanning Britain’s separation from continental Europe. We find overwhelming support for agriculture being introduced by incoming continental farmers, with small and geographically structured levels of additional hunter-gatherer introgression. We find genetic affinity between British and Iberian Neolithic populations indicating that British Neolithic people derived much of their ancestry from Anatolian farmers who originally followed the Mediterranean route of dispersal and likely entered Britain from northwestern mainland Europe.

---

The transition to farming marks one of the most important shifts in human evolution, impacting on subsistence, social organisation, health and disease vulnerabilities, economy, and material culture. The processes by which this transition occurred have been a matter of intense debate for over a century^10–15^, although across continental Europe recent ancient DNA studies indicate a predominant role for expanding Neolithic farmer populations of mostly Anatolian ancestry (Anatolian farmers - ANF)^1–9^. Anatolian farmer-derived populations dispersed throughout Europe via two major routes - one along the Mediterranean and the other through Central and into Northern Europe^3–7^. Both dispersals involved repeated, but mostly subsequent introgressions with local Mesolithic foragers, producing distinct cultural and genetic trajectories.

The nature of the Neolithic transition in Britain remains a puzzle because of the millennium-long delay in its appearance after the establishment of farming in adjacent regions of continental northwestern Europe^10–15^, and the lack of genome-wide data from British Mesolithic hunter-gatherers. Whilst there is universal agreement amongst archaeologists that there was a dramatic change in material culture in Britain after 6 kBP, there are divergent views regarding the extent to which this change was influenced by cultural or demographic processes^1–15^. One interpretation of the archaeological evidence is that British Mesolithic hunter-gatherers adopted Neolithic cultural practices abruptly at c.6 kBP without substantial immigration following prolonged contact with their continental neighbours^15^. This view is inconsistent with the predominantly Anatolian farmer-related ancestry in published data from British Early Neolithic farmers^8^, but the extent to which local British hunter-gatherer populations contributed to the first British farming populations, as well as the relationship of British hunter-gatherers to continental hunter-gatherer populations remains unresolved. These questions are of interest as Britain is situated between two genetically-distinct contemporaneous groups of Mesolithic hunter-gatherers – Western European and Scandinavian (WHGs & SHGs)^16^. The British Isles could also have putatively harboured ancestry from hunter-gatherers related to earlier Magdalenian Palaeolithic groups that recolonised Europe from the southwest after the Last Glacial Maximum (~21 to 17 kBP)^17^.

Here, we report the first genome data from six Mesolithic (including the ‘Cheddar Man’ skeleton from Gough’s Cave, Cheddar Gorge, Somerset) and 16 Neolithic British individuals, and combine it with new and already reported data from 51 previously published Neolithic British individuals^8^ to characterise the Mesolithic and Neolithic populations of Britain (Figure 1). We combined data generated in two different ways. For 35 individuals we generated new whole genome shotgun sequencing data (median coverage 0.09x) including the first full genomes from the British Mesolithic (at 2.3x) and Neolithic (at 10.7x). For all individuals we enriched next generation sequencing libraries for sequences overlapping about 1.24 million single nucleotide polymorphisms (SNPs) (median coverage 0.88x). We merged data obtained from both methods when it was available and chose the most likely base to represent the allele at each SNP (see Material and Methods). We merged the British Mesolithic and Neolithic data with 67 previously reported ancient DNA samples^1–3, 5–8, 10, 17–23^ (see Supplementary Table S1) and finally also with sequencing data from present-day individuals from diverse populations around the world^47^.

**Figure 1:**
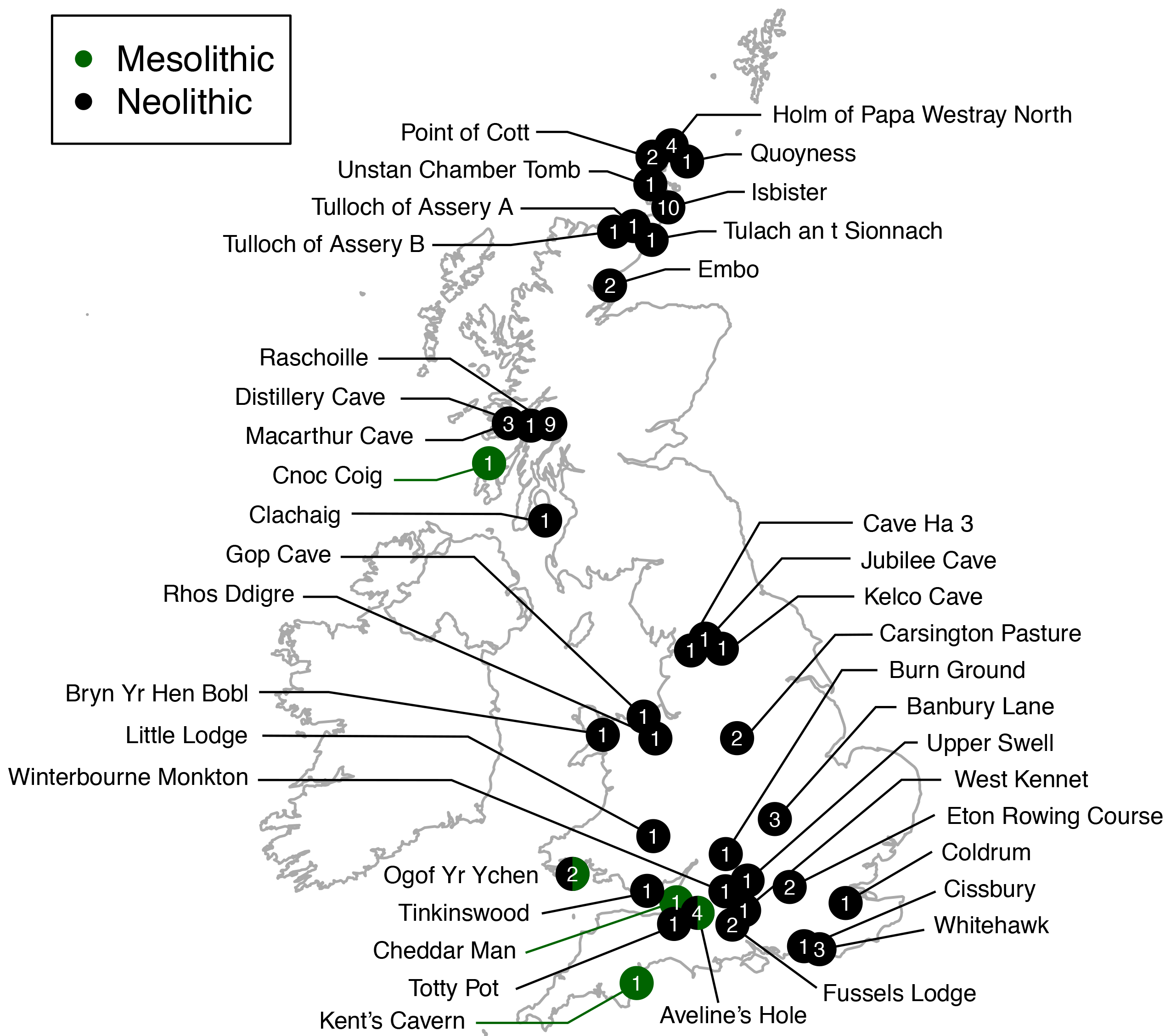
Map of sample locations. Geographical locations of British samples analysed here. Numbers indicate the number of samples from a given location.

We used principal components analysis to visualise some of the affinities of British Mesolithic and Neolithic genomes alongside those from ancient and modern West-Eurasian populations (Figure 2). The British Mesolithic individuals all cluster with Western and Scandinavian hunter-gatherers. By contrast, all directly-dated individuals who post-date 6 kBP, and undated individuals associated with Neolithic monuments, cluster tightly near Iberian and Central European Middle Neolithic individuals.

**Figure 2:**
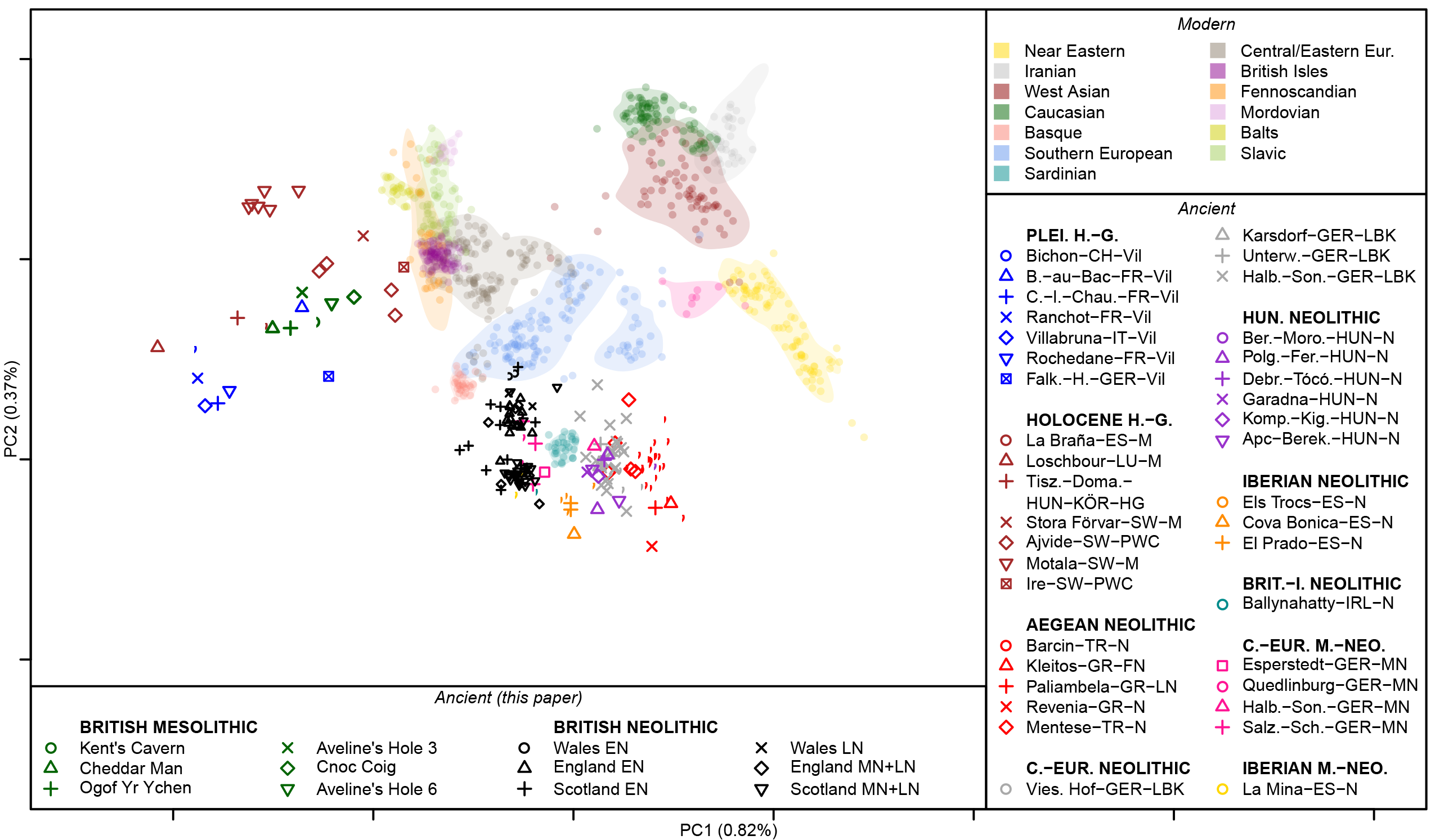
PCA of modern and ancient West-Eurasians. British and additional ancient samples are projected onto the reference space computed on modern West-Eurasian populations. See Material and Methods for computational details and Supplementary Table 1 for information on the samples. *Abbreviations:* European (Eur.), Pleistocene (Plei.), hunter-gatherer (H.-G.), British Isle (Brit.-I.), Middle Neolithic (M.-Neo.)

By examining the degree of allele sharing of British Mesolithic individuals to various European hunter-gatherer individuals or groups (SHG, EHG and El Miron, see Supplementary Figures S1-S4), we were able to attribute them confidently to the WHG. Comparison of British Mesolithic samples to different Mesolithic WHGs (Loschbour - Luxembourg, La Brana - Spain, KO1 - Hungary; Supplementary Figures S5-S6) indicates that they all resemble Loschbour most closely (i.e. the geographically most proximate Mesolithic genome available). When we compared the remaining British Mesolithic genomes to Loschbour and Cheddar Man (our highest-coverage British Mesolithic sample; ~2.3x), we found no significant excess of shared drift for either individual, indicating that Loschbour and the British Mesolithic samples do not form separate clusters (Supplementary Figure S7).

To investigate the proportions of Anatolian farmer-related ancestry in the British samples we modeled them as mixtures of ANFs and European WHGs (Figure 3, Supplementary Figure S8). All British Mesolithic samples could be explained entirely by WHG ancestry within the error bounds of the test. The majority (~75%) of ancestry in all British Neolithic individuals could be attributed to ANFs, indicating a substantial demographic shift with the transition to farming. These proportions of British Neolithic ANF/WHG ancestry are similar to Early Neolithic Iberian and Middle Neolithic Central European samples. We inferred some geographic structure in WHG admixture proportions among the British Early Neolithic samples; individuals from Wales retain the lowest levels of WHG admixture, followed by those from South-West and Central England. South East England and Scotland show the highest WHG admixture proportions. These proportions remain stable for over a millennium, from the Early into the Middle/Late Neolithic.

**Figure 3:**
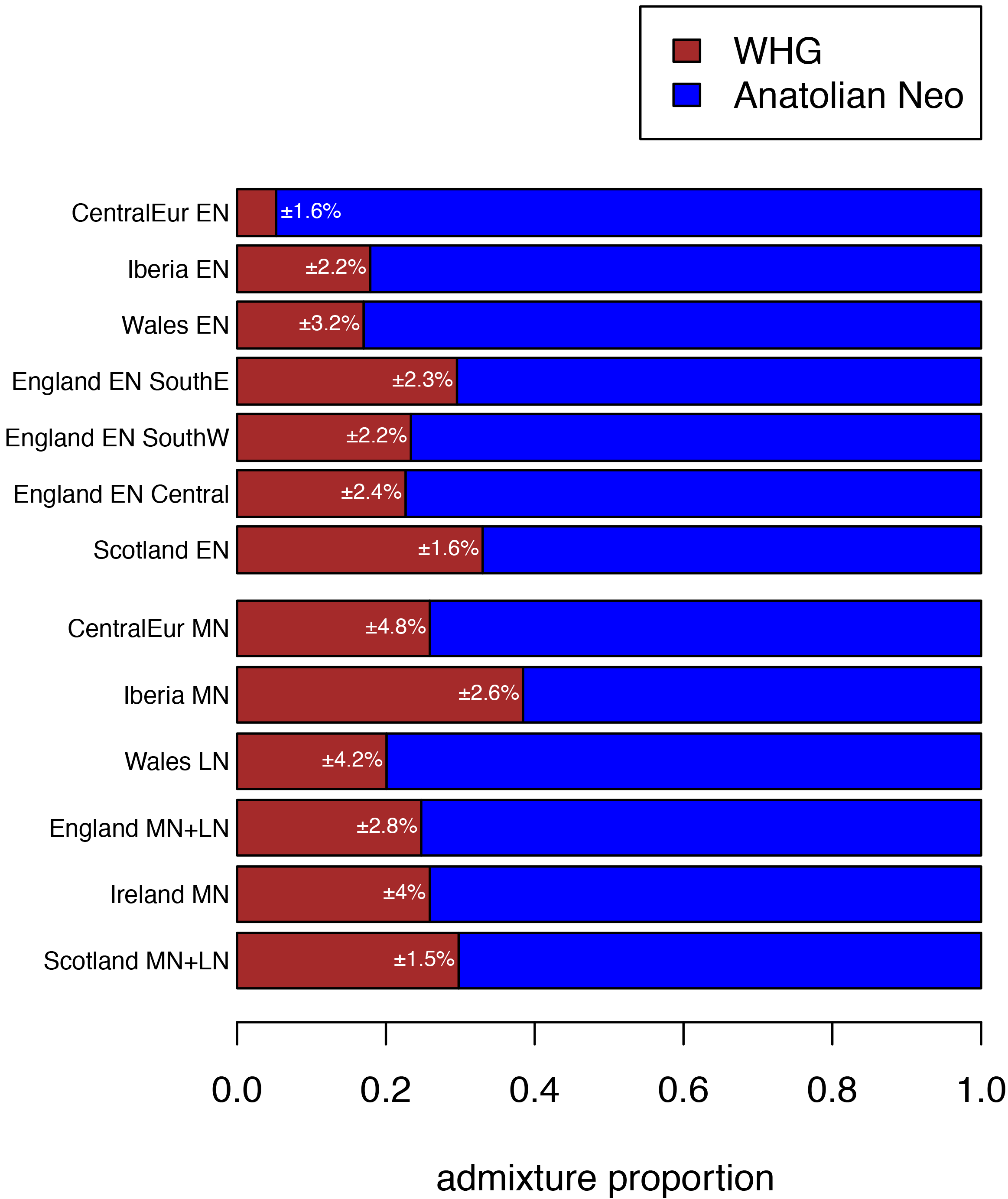
WHG and ANF ancestry components of British and Central European Neolithic populations. The relative WHG and ANF ancestry in Early and Middle Neolithic British and continental European populations quantified by *f*_*4*_ admixture proportions. Percentages indicate error estimates. See Material and Methods for computational details and Supplementary Table 1 for the lists of samples grouped into the different Neolithic populations. *Abbreviations:* Neolithic (Neo.), Early-, Middle-, and Late Neolithic (EN, MN, LN), South East (SouthE), South West (SouthW)

To investigate the proximate source of ANF ancestry in British Neolithic samples, we examined affinities with available Early Neolithic individuals from Iberia and Central Europe. We chose to compare Early over Middle Neolithic Iberians as the latter are contemporary with the British Early Neolithic, making them an unlikely direct source. For all of our British Neolithic individuals we inferred more shared drift with Early Neolithic Iberians; for the majority of comparisons this was significant (Figure 4A, Supplementary Figure S9). To infer levels of WHG introgression occurring between Iberian Early Neolithic populations – the closest currently available attributable source of farmer ancestry in British Early Neolithic genomes – and early British farmers, we estimated *f*_*4*_ admixture proportions. We detected little excess (~10%) WHG ancestry beyond what was already present in Iberian Early Neolithic populations, suggesting small proportions of admixture, particularly in Wales where we detected no excess WHG ancestry (Figure 4B). The small amounts of WHG introgression inferred here could have occurred on mainland Europe, and there is no need to invoke any genetic contribution from British Mesolithic hunter-gatherers to explain these results, although the significant bias towards British WHGs in some British Neolithic farmers, suggests that at least some of this introgression probably did occur in Britain (Supplementary Figure S7).

**Figure 4:**
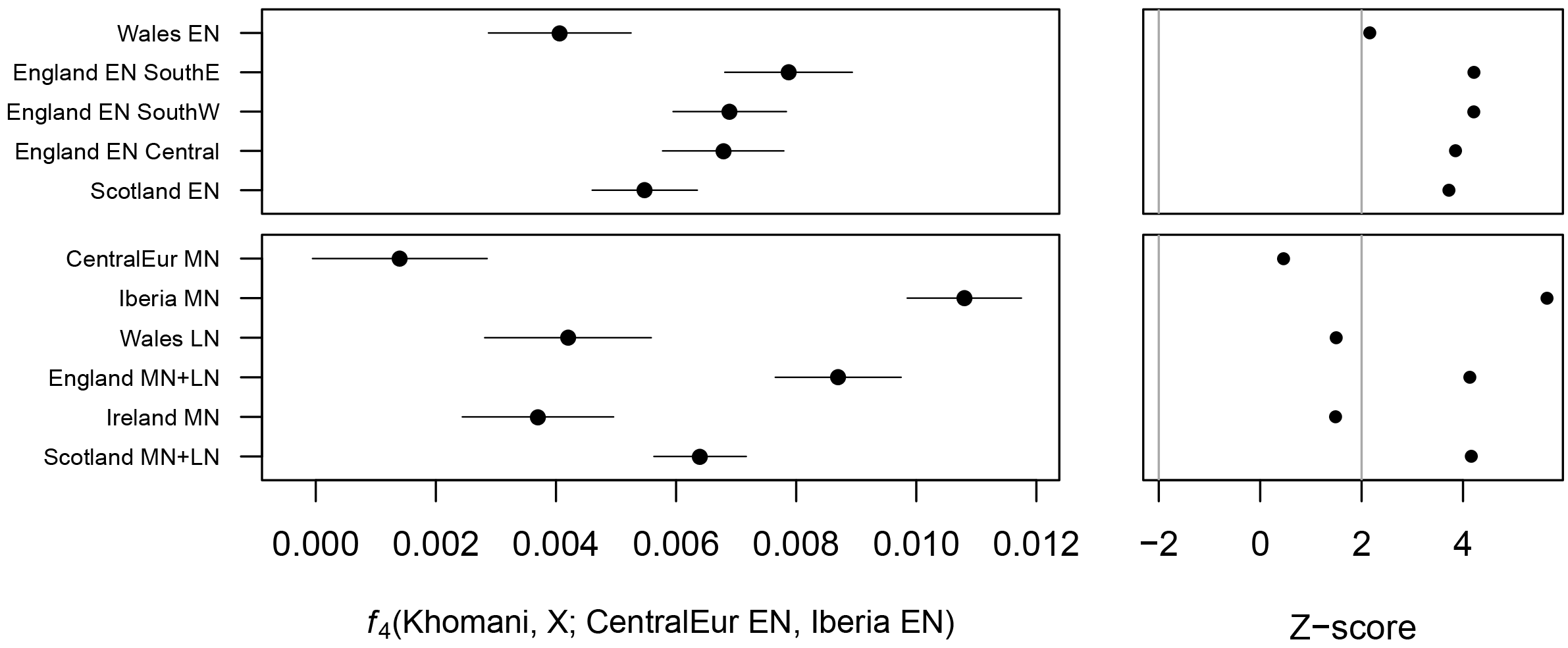
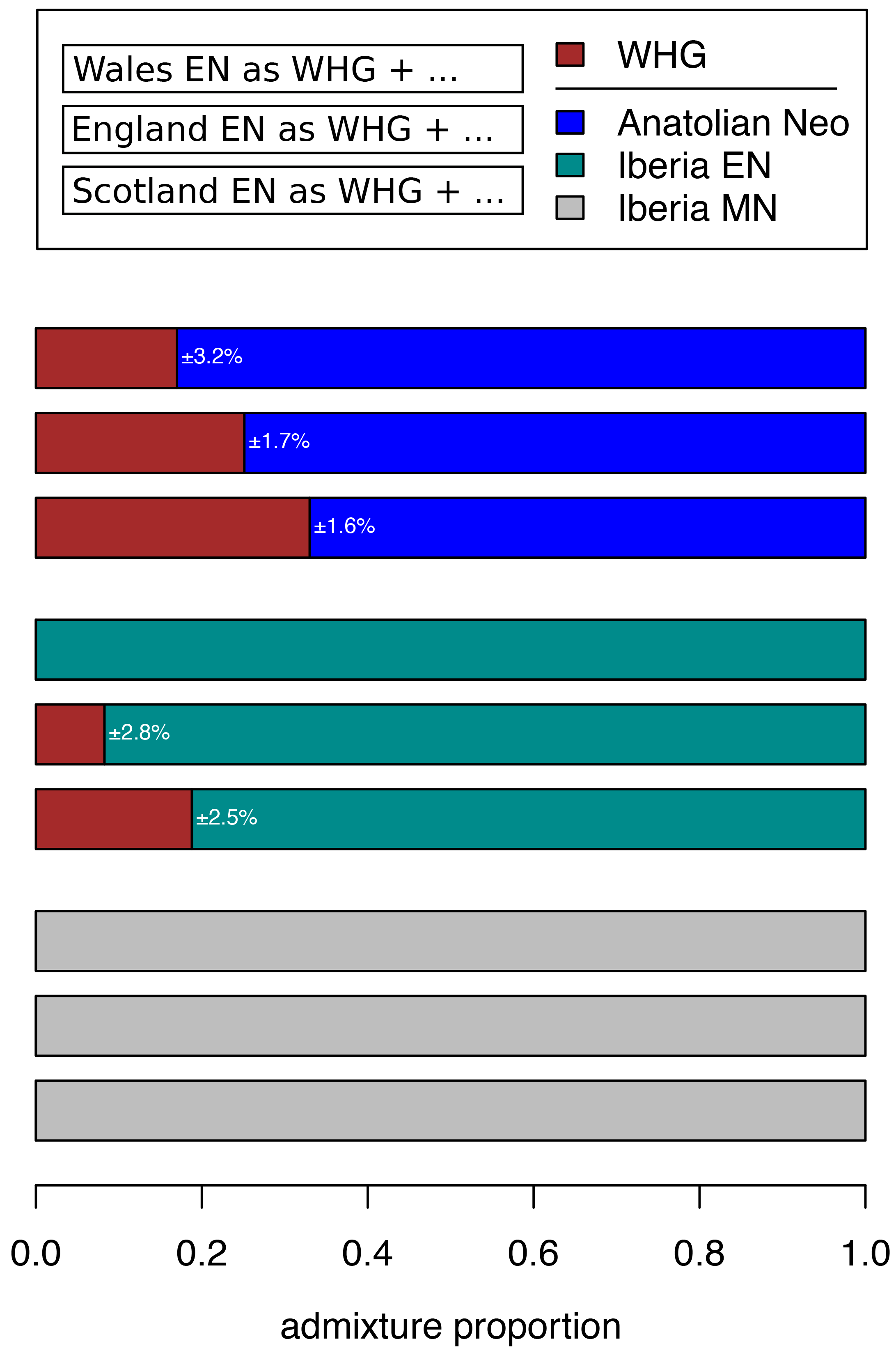
(A) Affinities of British and continental Neolithic populations. We compute admixture *f*_*4*_ statistics for different British EN, MN, and LN and continental MN populations comparing shared drift with Central European EN and Iberian EN populations. A positive Z-score above 2 corresponds to a significant affinity to the Iberian EN over Central European EN population. **(B)** *Quantifying excess WHG ancestry in British EN compared to Iberian EN and MN populations*. We compute *f*_*4*_ admixture proportions of WHG and different ANF populations in EN samples from Wales, England, and Scotland.

The Mesolithic Cheddar Man and the Neolithic sample from Carsington Pasture Cave, Derbyshire (‘Sven’) had sufficient coverage to estimate heterozygosity. Consistent with recent ancestry from larger or more admixed populations, Sven showed slightly higher levels of heterozygosity than Cheddar Man (Supplementary Figure S10). None of the Mesolithic and Neolithic British individuals analysed here had a derived lactase persistence allele (see Supplementary Table S2). We predicted pigmentation characteristics for Cheddar Man and Sven using Hirisplex^25^ and a recently-developed method for predicting skin pigmentation^26^. Previously, predictions on the level of skin pigmentation were mostly derived using two SNPs in SLC45A2 and SLC24A5 that indicate lack of hypo-pigmentation when in the ancestral state^18^. However, here we integrate 36 rather than 2 SNPs allowing more precise prediction^26^. Cheddar Man is predicted to have had dark or dark to black skin, blue/green eyes and dark brown possibly black hair, whereas Sven most likely had intermediate to dark skin pigmentation, brown eyes and black possibly dark brown hair (see Pigmentation section in the Supplementary Materials for a detailed discussion of the results). This is in line with the current hypothesis that alleles commonly associated with lighter skin were introduced in Western Europe by ANFs^19^.

We also analysed two previously-published WHGs, and find potential temporal and/or geographical variation in pigmentation characteristics. Loschbour^22^ from Luxembourg is ~2000 years younger than Cheddar Man, and is predicted to have had intermediate skin pigmentation. Furthermore, the Loschbour individual most likely had blue/green eyes. In contrast, La Braña^18^ from northern Spain who is slightly later than Loschbour is predicted to have had dark to dark to black skin and hazel/green eye colour. Both La Braña and Loschbour were predicted to have had black, possibly dark brown hair. These results imply that quite different skin pigmentation levels coexisted in WHGs at least by around 6000 BC.

## Discussion

The genomes of the six British Mesolithic individuals examined here appear to be typical of WHGs, indicating that this population spread to the furthest northwestern point of early Holocene Europe after moving from southeastern Europe, or further east, from approximately 14,000 years ago^17^. It is notable that this genetic similarity among British Mesolithic and Loschbour individuals spans a period in Britain (c.10.5-6 kBP) that includes the cultural transition to the Late Mesolithic, the potentially catastrophic Storegga landslide tsunami, and the separation of Britain from continental Europe as sea levels rose after the last ice age. This finding is inconsistent with the hypothesis of pre-c.6 kBP gene flow into Britain from neighbouring farmers in continental Europe^15^.

Our analyses indicate that the appearance of Neolithic practices and domesticates in Britain c.6 kBP was mediated overwhelmingly by immigration of farmers from continental Europe^2,13–14^, and strongly reject the hypothesised adoption of farming by indigenous hunter-gatherers as the main process^15^. British farmers were substantially descended from Iberian Neolithic-related populations whose ancestors had expanded along a Mediterranean route^2,7,14^, although our analyses cannot rule out the possibility that they also inherited a minority portion of their ancestry from the Danubian route expansion through Central Europe. Indeed, a recent study that investigated continental Neolithic farmer-related ancestry components in Neolithic Britain estimated ⅔ Mediterranean and ⅓ Danubian route^8^, which may be consistent with the association between Britain’s more easterly-distributed Carinated Bowl tradition^14^ and the Nord-Pas-de-Calais region of France, as Neolithic people in these regions of mainland Europe are thought to have interacted with populations of Central European Neolithic ancestry^14, 27–29^. The varied Neolithic cultures of northern France, Belgium and the Netherlands, the most likely continental sources for the British Neolithic^14^, may represent a genetically heterogeneous population who shared large but variable proportions of their ancestry with Neolithic groups in Iberia via Atlantic and southern France.

We caution that our results should not be interpreted as showing the Iberian Neolithic-related ancestry in British Neolithic people derives from migrants whose ancestors lived in Iberia, as we do not have ancient DNA data from yet unidentified source populations — possibly located in southern France — that were ancestral to both Iberian and British farmers. Available Middle Neolithic Iberian individuals are too late to represent the source population for early British farmers, and there is no archaeological evidence for direct immigration from Iberia^14^. The lack of genome-wide data from Neolithic northern France, Belgium and the Netherlands means that it is not currently possible to identify proximal continental source populations.

The limited regional variation in WHG ancestry we see in the British Neolithic samples could reflect subtle but differing degrees of regional admixture between farmers and foragers, and/or multiple continental source populations carrying varying levels of WHG ancestry colonising different regions of Britain. What is clear is that across Britain all of our estimates for admixture between hunter-gatherers and farmers are very small, and that we find no evidence of WHG ancestry increasing as the British Neolithic progressed over time (Figure 3). In contrast, the resurgence of WHG ancestry in all available continental European Middle Neolithic samples, prior to the British Neolithic transition and including a population sample from southern France, means that the level of WHG ancestry we see in most British Neolithic farmers could be accounted for entirely in continental source populations^3, 8–10^.

Evidence for only low levels of WHG introgression among British Neolithic people is striking given the extensive and complex admixture processes inferred for continental Neolithic populations^3^’ ^8–10, 30–32^. Low levels of admixture between these two groups on the wavefront of farming advance in continental Europe have been attributed to the groups maintaining cultural and genetic boundaries for several centuries after initial contact^30–32^. Similarly, isotopic and genetic data from the west coast of Scotland are consistent with the coexistence of genetically distinct hunter-fisher-gatherers and farmers, albeit for a maximum of few centuries ^33–34^. The resurgence in WHG ancestry after the initial phases of the Neolithic transition in continental Europe indicates that the two populations eventually mixed more extensively^3, 8–10^. However there is no evidence for a WHG resurgence in the British Neolithic up to the Chalcolithic population movements associated with the Beaker phenomenon (c.4.5 kBP)^8^. This is consistent with the lack of evidence for Mesolithic cultural artefacts in Britain much beyond 6 kBP^14^ and with a major dietary shift at this time from marine to terrestrial resources; particularly apparent along the British Atlantic coast^33–35^.

In summary, our results indicate that the progression of the Neolithic in Britain was unusual when compared to other previously studied European regions. Rather than reflecting the slow admixture processes that occurred between ANFs and local hunter-gatherer groups in areas of continental Europe, we infer a British Neolithic proceeding with little introgression from resident foragers – either during initial colonization phase, or throughout the Neolithic. This may reflect the fact that farming arrived in Britain a couple of thousand years later than it did in Europe. The farming population who arrived in Britain may have mastered more of the technologies needed to thrive in northern and western Europe than the farmers who had first expanded into these areas. A large-scale seaborne movement of established Neolithic groups leading to the rapid establishment of the first agrarian and pastoral economies across Britain, provides a plausible scenario for the scale of genetic and cultural change in Britain.

## Materials and Methods

### Ancient DNA Extraction and Sequencing

DNA extractions and library preparations for all samples with newly reported data were conducted in a dedicated ancient DNA laboratory (NHM, London). We used approximately 25mg of finely drilled bone powder and followed the DNA extraction protocol described in Dabney *et al.*^36^ but replaced the Zymo-Spin V column binding apparatus with a high pure extender assembly from the High Pure Viral Nucleic Acid Large Volume Kit (Roche). Library preparations followed the partial uracil-DNA-glycosylase treatment described in Rohland *et al.*^37^ and a modified version of the Meyer & Kircher^38^ protocol. Library modifications: the initial DNA fragmentation step was not required; all clean-up steps used MinElute PCR purification kits (Qiagen). The index PCR step included double indexing^39^, the polymerase AmpliTaq Gold and the addition of 0.4mg/mL BSA. The index PCR was set for 20 cycles with three PCR reactions conducted per library. Libraries were screened for DNA preservation on an Illumina NextSeq platform, with paired-ends reads. Promising libraries were further enriched in two ways, one at the NHM using in-solution hybridisation capture enrichment kits (Mybaits-3) from MYcroarray. The baits were designed to cover ca. 20K SNP’s (5,139 functional and 15,002 neutral SNP’s) at 4х tiling. Capture protocol followed the manufacturers instructions in the Mybaits manual v3. Captured libraries were sequenced on an Illumina NextSeq platform (NHM) using paired-ends reads. Newly reported data from 36 of these libraries was also obtained at the dedicated ancient DNA lab in Harvard Medical School by enriching in solution for approximately 1.24 million targeted SNPs. We sequenced these libraries on an Illumina NextSeq500 instrument, iteratively sequencing more until we estimated that the additional number of targeted SNPs hit per newly generated sequence was less than 1 per 100.

### Bioinformatics

All sequence reads underwent adapter and low-quality base trimming, and overlapping reads pairs were collapsed with AdapterRemoval^40^. Non-collapsed reads and those below 30bp were discarded, and the remaining aligned against the hs37d5 human reference genome with BWA^41^. Mapped reads with MAPQ at least 30 were merged per individual and re-aligned around InDels with GATK^42^. Resulting BAM files were split by flowcell and lane, and empirical ATLAS^43^ post mortem damage patterns estimated per individual per lane for lanes with at least 5.5 million reads, otherwise per individual per flowcell. ATLAS BQSR base quality score recalibration tables were generated per lane for lanes with at least 5.5 million reads, otherwise per flowcell. We generated recalibrated BAM files per individual with ATLAS recalBAM, and used those to estimate mitochondrial contamination and determine Mitochondrial haplogroups with ContamMix^44^ and Phy-Mer^45^ respectively. We considered Mitochondrial contamination to be tolerable if 0.98 was included in the confidence intervals. Haploid genotypes were called with ATLAS allelePresence with theta fixed at 0.001, determining the most likely base at a position. Heterozygosity estimates shown in Sup. Fig. S10 were computed with ATLAS estimateTheta and default window size of 1Mbp, excluding windows that overlap with telo- or centromeres.

### PCA

Principal component analysis was performed with LASER^46^ following the approach described previously^5^. After generating a reference space of modern Western Eurasian individuals^6^, we projected the BAM files of ancient reference individuals (see Supplementary Table S1 for references) and the British individuals presented here into the reference space via Procrustes analysis implemented in LASER.

### f-statistics

The various flavours of *f*-statistics presented here, *i.e*. outgroup *f*_*3*_, *f*_*4*_, and *f*_*4*_ admixture proportions, were computed with *qpPop, qpDstat* in *f*_*4*_ mode, and *qpAdm* from the ADMIXTOOLS^24^ package with default parameters on the positions defined by the *HOIll* set of SNPs^6^. Ancient individuals analysed here are listed in Sup. Table 1. Modern reference individuals were first published in Mallick *et al.*^47^. All *qpAdm* runs used the set of outgroups Han, Karitiana, Mbuti, Onge, Papuan, Mota, Ust_Ishim, MA1, ElMiron, GoyetQ116-1.

## Acknowledgements

The authors are extremely grateful to individuals and institutions who granted permissions to sample human remains, gave their time to facilitate sampling and lent their invaluable expertise on collections: the Longleat Estate, Tom Lord at Lower Winskill Farm, Barry Chandler at Torquay Museum, Andrew Chamberlain at the University of Manchester, Linda Wilson and Graham Mullan at the University of Bristol Spelaeological Society, Elizabeth Walker, Adam Gwilt and Jody Deacon an at the National Museum of Wales, Andy Maxted at Brighton Museum, Marta Lahr at the Duckworth Laboratory, Barry Lane at Wells Museum, Martin Smith at Bournemouth University, David Rice at the Museum of Gloucester and Rob Kruszinski at the Natural History Museum. In addition, Y.D. wishes to thank Jens Blöcher, Amelie Scheu, Christian Sell and Joachim Burger for the many valuable discussions on the bioinformatic pipeline, and Vivian Link for help with ATLAS. D.R. was supported by NIH grant GM100233, by NSF HOMINID BCS-1032255, and by an Allen Discovery Center of the Paul Allen Foundation, and is a Howard Hughes Medical Institute investigator. M.G.T. and I.B. were supported by a Wellcome Trust Investigator Award (project 100713/Z/12/Z).

